# Spatial predictive coding in visual cortical neurons

**DOI:** 10.1101/2025.09.17.676794

**Authors:** Qingqing Zhang, Sverre Grødem, Alexa Gracias, Kristian K. Lensjø, Marianne Fyhn, Carsen Stringer, Marius Pachitariu

## Abstract

Predictive coding is a theoretical framework that can explain how animals build internal models of their sensory environments by predicting sensory inputs. Predictive coding may capture either spatial or temporal relationships between sensory objects. While the original theory by Rao and Ballard, 1999 described spatial predictive coding, much of the recent experimental data has been interpreted as evidence for temporal predictive coding. Here we directly tested whether the “mismatch” neural responses in sensory cortex are due to a spatial or a temporal internal model. We adopted two common paradigms to study predictive coding: one based on virtual-reality and one based on static images. After training mice with repeated visual stimulation for several days, we performed multiple manipulations, including: 1) we introduced a novel stimulus, 2) we replaced a stimulus with a novel gray wall, 3) we duplicated a trained stimulus, or 4) we altered the order of the stimuli. The first two manipulations induced a substantial mismatch response in neural populations of up to 20,000 neurons recorded across primary and higher-order visual cortex, while the third and fourth ones did not. Thus, a mismatch response only occurred if a new spatial – not temporal – pattern was introduced.

## Introduction

Predictive coding has emerged as a popular all-encompassing neural theory which postulates that neural representations in sensory brain areas are optimized to predict sensory inputs [1–3]. To do this prediction well, neuronal networks must contain an intrinsic model of the natural world which can be used to detect unpredicted changes in the environment. Predictive coding theories can be divided into temporal and spatial predictive coding. Temporal predictive coding is the most prevalent form, and often implied implicitly [4–6]. It postulates that brains are optimized to predict future sensory inputs from past sensory inputs and/or from self-movements. Spatial predictive coding, while less popular as a general theory of brain function, was the basis of the original predictive coding framework [7, 8]. It postulates that brains predict localized sensory inputs from their larger spatial context, with no requirement for temporal information.

Predictive coding is a learning theory with substantial computational capabilities. Some of the most highly-capable artificial systems are trained using cost functions that are similar predictive coding [9]. These artificial intelligence (AI) methods vary in how they treat the temporal and spatial domains. For example, large language models are trained to predict the next word in a sequence, a temporal prediction [10]. In contrast, large vision models are often trained to predict left-out patches in images from surrounding patches, a spatial prediction [11, 12]. Vision models for video analysis are pretrained using a combination of spatial and temporal prediction [13, 14]. While the cost functions for temporal and spatial predictive coding share similar principles, their underlying mechanistic implementations and intended functions differ substantially. Thus, it is important to distinguish between different types of predictive coding, in order to understand the inner workings of a system like the mammalian sensory cortex.

The main evidence for predictive coding in the brain has come from observations of the “mismatch response” or sensory “prediction error” in sensory cortex [5, 6, 15]. These are large neural responses that are observed under sensory manipulations that deviate from the expected, or typical sensory inputs. In most cases, the expectations are established spatially, temporally, and/or by the novelty of a stimulus [16]. Such large neural responses are observed in both humans and animals, using a variety of recording methods from single-unit recordings [4, 17–22], to LFP [23–26], fMRI [27–31], and EEG/MEG [32–36]. Focusing on single-unit recordings in animals, the mismatch response has been observed in response to sensory manipulations after a specific stimulus sequence had been shown repeatedly over many trials and across several consecutive days [6, 15]. In some studies, the stimulus sequence was coupled to motor drive, for example in closed-loop virtual reality where mice control the forward progression through a corridor [37, 38]. The motor coupling is sometimes thought to be critical, as it could enable animals to predict the consequences of their own actions [39–42].

Here we perform a critical evaluation of the predictions of spatial and temporal predictive coding, using two closed-loop sensory paradigms designed to be similar to previous studies. We recorded large-scale neural populations from primary and higher-order visual cortices. Using various sensory manipulations with appropriate controls, we find that the mismatch response in primary and higher-order visual areas represents a spatial, rather than a temporal prediction error.

## Results

In our first set of experiments, mice ran for five days in a virtual reality environment in single sessions of up to 1.5 hours (Figure 1A). The mice ran forward through linear corridors at an average speed of 19.4cm/s, experiencing in each trial a succession of four visual landmarks. This was similar to [6], except we used crops from natural images rather than gratings for increased saliency and ecological relevance. We denote individual stimuli by capitalized letters, followed by a subscripted number indicating the position of the landmark in the trial. Thus, standard trials are denoted A_1_B_2_C_3_D_4_, while manipulated test trials during test sessions are A_1_B_2_X_3_D_4_, with X = N, D etc. In-between landmarks, the corridors had a fixed pattern of sparse noise squares which never changed. All individual visual landmarks (i.e. C_3_) were composed of a pair of repeated sub-images (i.e. cc). The role of these images was to allow the introduction of half-landmarks (i.e. c) or repeated landmarks (i.e. ccc) without introducing new spatial patterns locally (see next section).

**Figure 1:**
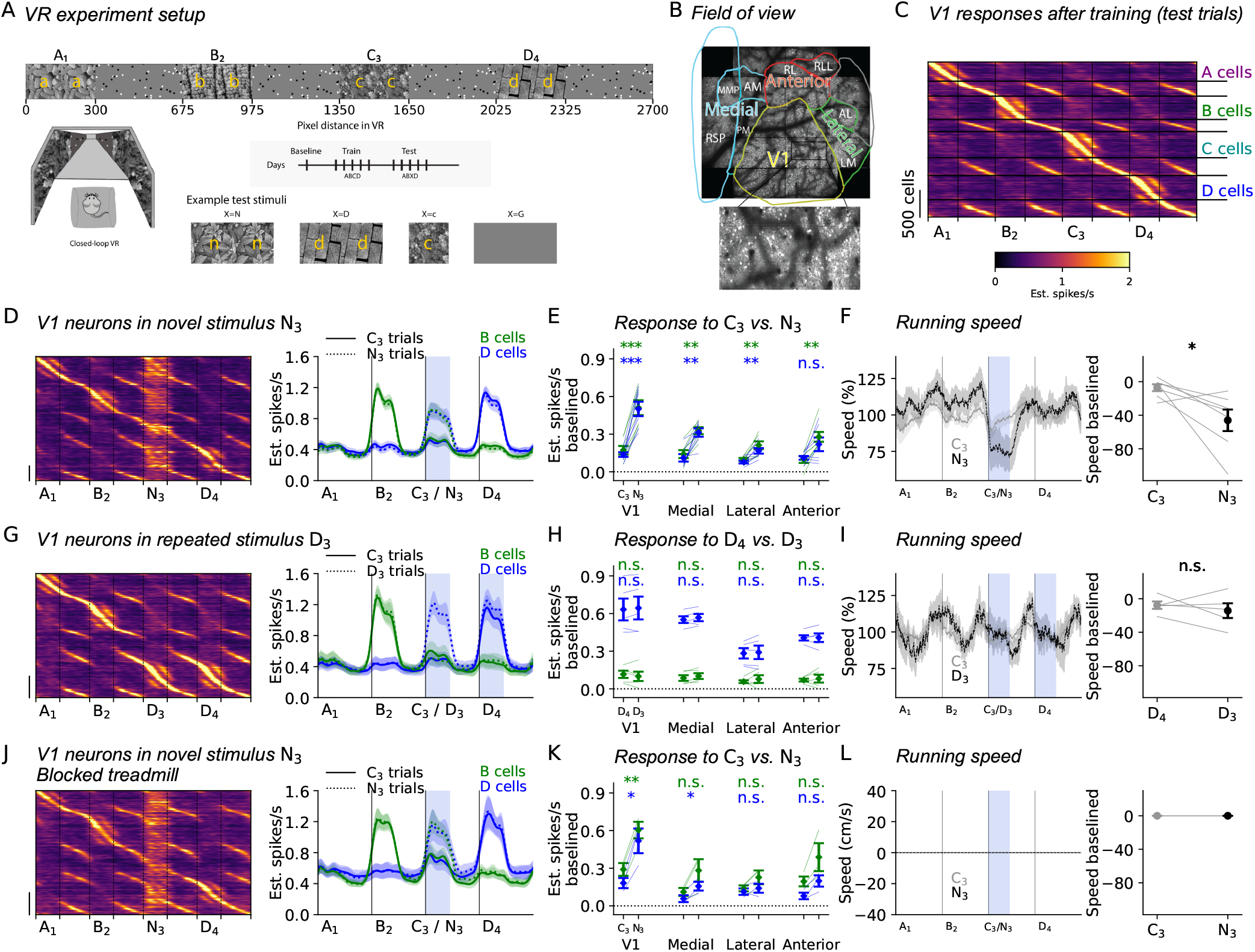
Mismatch neural responses to novel sensory inputs. (A) Experimental setup illustrating the progression through the VR corridor (top), the 270 degree virtual reality setup (bottom left) and the progression of training and testing sessions over days. (B) Field of view of the neural recordings with zoom-in. (C) Trial-averaged neural responses after training. Neurons are sorted by preferred spatial position and responses are shown on held-out trials. (D) (left) Same as (C) for trials where a new stimulus N_3_ was introduced. Preferred spatial positions were computed on A_1_B_2_C_3_D_4_ trials excluding the C_3_ positions; (right) Population-averaged responses (n=7 mice). (E) Average responses from (D) in the shaded blue area. (F) (left) Average running speeds for A_1_B_2_C_3_D_4_ and A_1_B_2_N_3_D_4_ trials, and (right) quantification in the blue shaded area. (G-I) Same as (D-F) for A_1_B_2_D_3_D_4_ trials (n=5 mice). (J-L) Same as (D-F) for open-loop A_1_B_2_C_3_D_4_ trials where the treadmill was blocked (n=3 mice). All data are mean *±* s.e.m. across mice. Statistical significance in all figures was assessed using paired two-sided t-tests across mice with *, **, and *** representing *p* < 0.05, *p* < 0.01, and *p* < 0.001 respectively.

During test sessions after training, we recorded neural activity simultaneously from populations of 5,846 - 22,711 neurons from primary visual cortex as well as higher-order visual areas using a two-photon mesoscope (Figure 1B). Neural activity was deconvolved and interpolated to the position of the corridor, then averaged over trials. Neurons were sorted according to their position tuning in the A_1_B_2_C_3_D_4_ corridor, and their averaged activity was computed on held-out trials, that were not used to calculate the peak tuning position (Figure 1C). Finally, neurons were assigned into groups based on the location of their peak firing (A, B, C, D cells in Figure 1C). Note that on some subsequent analyses, A, B, D cells were defined without including the C_3_ corridor portion, because that portion was subject to manipulations and quantifications (see Methods). We also chose to focus on B and D cells exclusively, since these have preferred responses close to the manipulated C_3_ region and would be more likely to show effects.

### Responses to novel spatial and temporal patterns

We first tested whether a novel spatial pattern N3 would drive a mismatch response, defined as a substantially larger neural response compared to the trained stimuli (Figure 1D). Both B and D cells responded substantially more to the novel stimulus N_3_ compared to the old pattern C_3_, nearly matching their firing rates to their own preferred stimuli (B_2_ and D_4_ respectively, Figure 1DE). This increased response was accompanied by a reduction in running speed in nearly all mice, as well as modest pupil dilation (Figure 1F, Figure S1A). The increased response was not present in anesthesia (Figure S2). The novel spatial pattern may thus have driven a mismatch response, either due to a failed prediction of stimulus C_3_ (temporal predictive coding), or due to the novelty of the spatial pattern in stimulus N_3_ (spatial predictive coding).

To disentangle these two hypotheses, we next tested whether replacing C_3_ with a trained pattern D_3_ also results in a mismatch response. If temporal predictive coding applies, we would still expect a failed prediction of stimulus C_3_. If mismatch responses instead result from spatial predictive coding, they should not occur here, because the spatial patterns in D_3_ also occur in D_4_, a trained stimulus. We observed no mismatch response on A_1_B_2_D_3_D_4_ trials, with neural responses to D_3_ being indistinguishable from those to D_4_ (Figure 1GH). We also did not observe changes in mouse behaviors such as running speed or pupil fluctuations (Figure 1I, Figure S1B).

Since mice did not indicate behaviorally that a “failed prediction” occurred for the D_3_ stimulus, it is possible that the mismatch responses to N_3_ are in fact directly due to behavioral modulations, such as decreased running speeds, which did occur in A_1_B_2_N_3_D_4_ trials (Figure 1F). To test this hypothesis, we performed test sessions with N_3_ trials in which we clamped the treadmill on all trials, thereby preventing the mice from running, while automatically moving the VR forward in open-loop. Similar to unclamped, closed-loop trials, we observed mismatch responses to N_3_ in the B and D cells that were larger than the responses to C_3_ and nearly as large as the responses to their preferred stimuli (B and D, respectively, Figure 1JK). Thus, mismatch responses were observed even without running (Figure 1L).

In our next test, we shortened the stimulus C_3_ to just its first half “c” (Figure 1A). No new spatial patterns were introduced in this way, since the spatial transitions from sparse noise to “c” and from “c” to sparse noise were already contained in the trained stimulus containing C_3_ =“cc” (Figure 1A). However, a new temporal pattern was introduced this way, which could lead to a failed prediction of a second “c”, should the temporal theory hold. We did not find a mismatch response to the missing second “c” (Figure 2AB), again arguing against the temporal predictive coding hypothesis. We also did not observe changes in running and pupil fluctuations (Figure 2C, Figure S1C).

**Figure 2:**
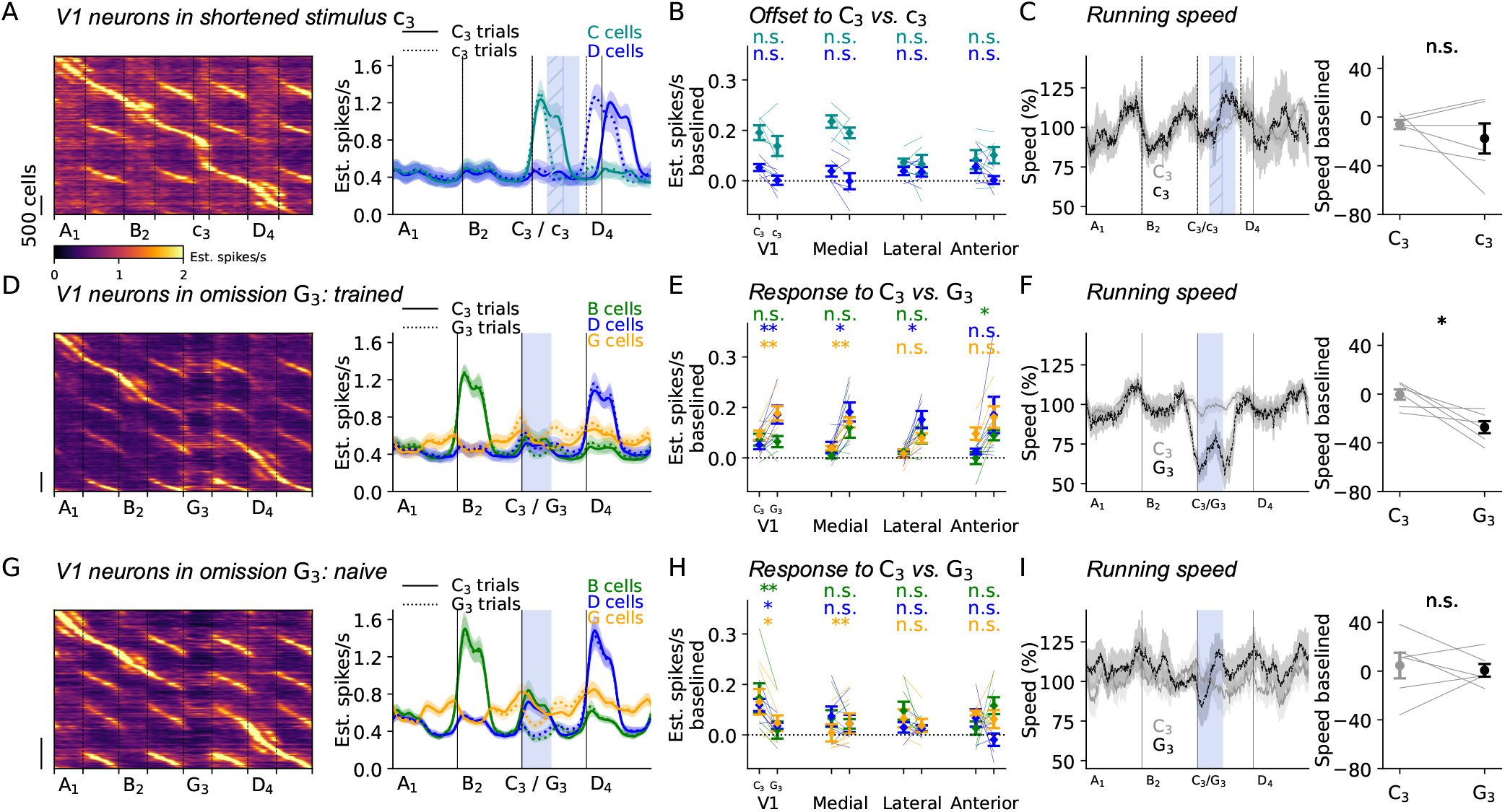
Additional manipulations and omissions for test stimuli. (A) Same as Figure 1D-F for a shortened stimulus manipulation (c_3_ = half of C_3_, n=6 mice). (D-F) Same as Figure 1D-F for trials where C_3_ is omitted and replaced with gray screen G_3_ (n=6 mice). (G-I) Same as (D-F), but before training in naive animals (n=6 mice). All data are mean*±*s.e.m. across mice. Paired two-sided t-tests were performed across mice.

### Understanding omission responses

In the next test we adopted a paradigm from previous studies [6], by omitting a trained stimulus C_3_ from the sequence and replacing it with a gray wall G_3_ (Figure 2D). Similar to previous studies, this omission resulted in elevated neural responses, which have previously been interpreted as a temporal mismatch error (Figure 2DE). In our data, these omission responses were substantially smaller than the responses to the novel stimulus N_3_ (compare to Figure 1DE). We also observed a behavioral response, similar to the N_3_ trials (Figure 2F). The elevated neural responses and behavioral changes to the gray wall were not observed in naive mice (Figure 2G-I). In previous studies, such findings were interpreted to suggest that omission responses are due to temporal prediction errors [6]. However, notice that the gray wall also introduces novel spatial patterns.

Before addressing omission responses further, we introduce another set of experiments, designed to be similar to other paradigms where temporal predictive coding was invoked, and where omission mismatch responses were similarly observed [5]. In these experiments, we presented sequences of static images A_1_B_2_C_3_G_4_ where G is a gray screen (Figure 3A). To maintain the closed-loop nature of the experiment, which is thought to be required for temporal predictive coding [40], we linked the image transitions to the running speed of the mice (averaging 29.8 cm/s), requiring mice to run for a set period before transitioning to the next stimulus. We used different durations for each stimulus to encourage temporal structure learning (2s, 1s, 3s and 1s respectively for A_1_, B_2_, C_3_ and G_4_). This created an abstract linear “corridor” in each trial, that we used to interpolate neural responses to, similar to the VR corridor. To focus on the fast neural responses to stimulus transitions, we recorded at faster frame rates (30Hz) and focused on individual recording locations in the primary visual cortex (Figure 3A). After training, neural responses were primarily concentrated at stimulus transitions, including at gray screen onset (Figure 3B).

**Figure 3:**
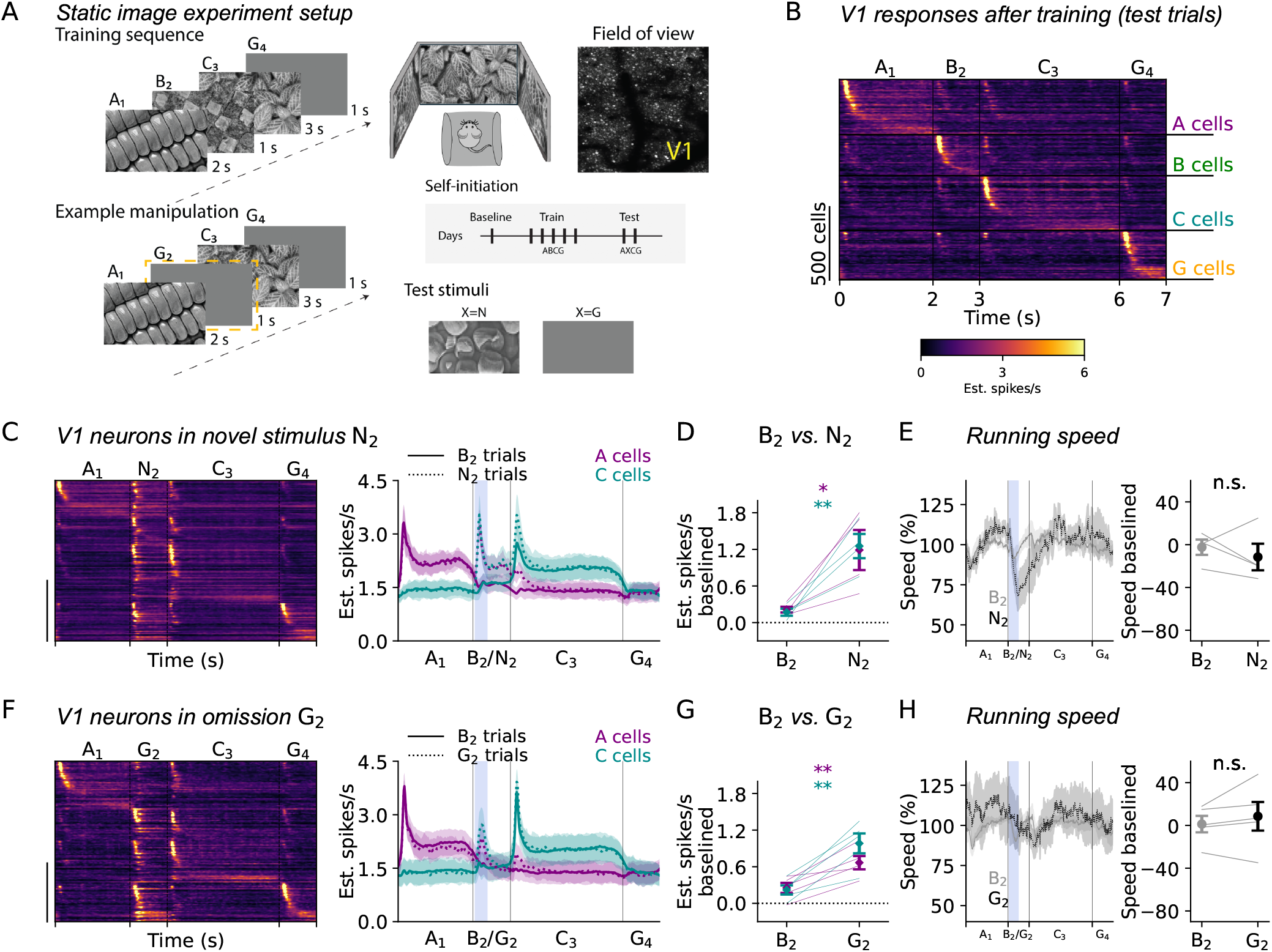
Mismatch responses with self-initiated static stimuli. (A) Experiment setup, similar to the VR experiments, but the forward running initiates transitions between static images, and the recorded field of view is smaller and focused on V1. Transitions between different stimulus pairs require different durations of constant forward running as shown in left panels. (B) Trial-averaged neural responses after training. Neurons were sorted by their peak response times and held-out trials are shown. (C) (left) Same as (B) for trials where a novel stimulus N_2_ was introduced. Cell sorting was done without the B_2_ stimulus. (right) Population-average for A cells and C cells (n=4 mice). (D) Quantification of baselined average responses inside the blue window in (C). (E) (left) Average running speeds for A_1_B_2_C_3_D_4_ and A_1_B_2_N_3_D_4_ trials, and (right) quantification in the blue shaded area. (F-H) Same as (C-E) for trials where B_2_ is omitted and replaced with a gray screen G_2_ (n=5 mice). All data are mean*±*s.e.m. across mice. Paired two-sided t-tests were performed across mice.

Similar to the VR experiments, we found that a novel stimulus N_2_ introduces elevated firing rates in multiple neural populations (Figure 3CD), although without a significant behavioral change (Figure 3E). Also similar to the VR experiments, “omission” trials where we replaced B_2_ with a gray screen *G*_2_ also resulted in elevated firing rates, potentially indicating mismatch errors (Figure 3FG). These were also not accompanied by clear behavioral changes (Figure 3H).

Thus, omission trial responses in both the VR and static image experiments appear to suggest temporal prediction errors. To illustrate why this is not necessarily the case, consider the bottom-up receptive field (RF) responses of visual cortical neurons (Figure 4A). The virtual corridors drive spatial RFs at every position, including at the boundary between the sparse noise and the gray screen (omitted C_3_). In this case, G_3_ introduces a new spatial pattern at this boundary, which would drive different sets of neurons based on their spatial RFs. Similarly, at image transitions in the static image experiment, spatiotemporal RFs (stRF) need to take into consideration the transition between one spatial pattern and another, since neurons as early as the retina respond to step-changes in illumination in an on/off fashion rather than to sustained illumination patterns [43]. A new stRF pattern is introduced at the transition between A_1_ and G_2_, that was not present in the trained stimuli. Thus, mismatch responses on omission trials can be explained by novel bottom-up inputs or novel spatial patterns (Figure 4A).

**Figure 4:**
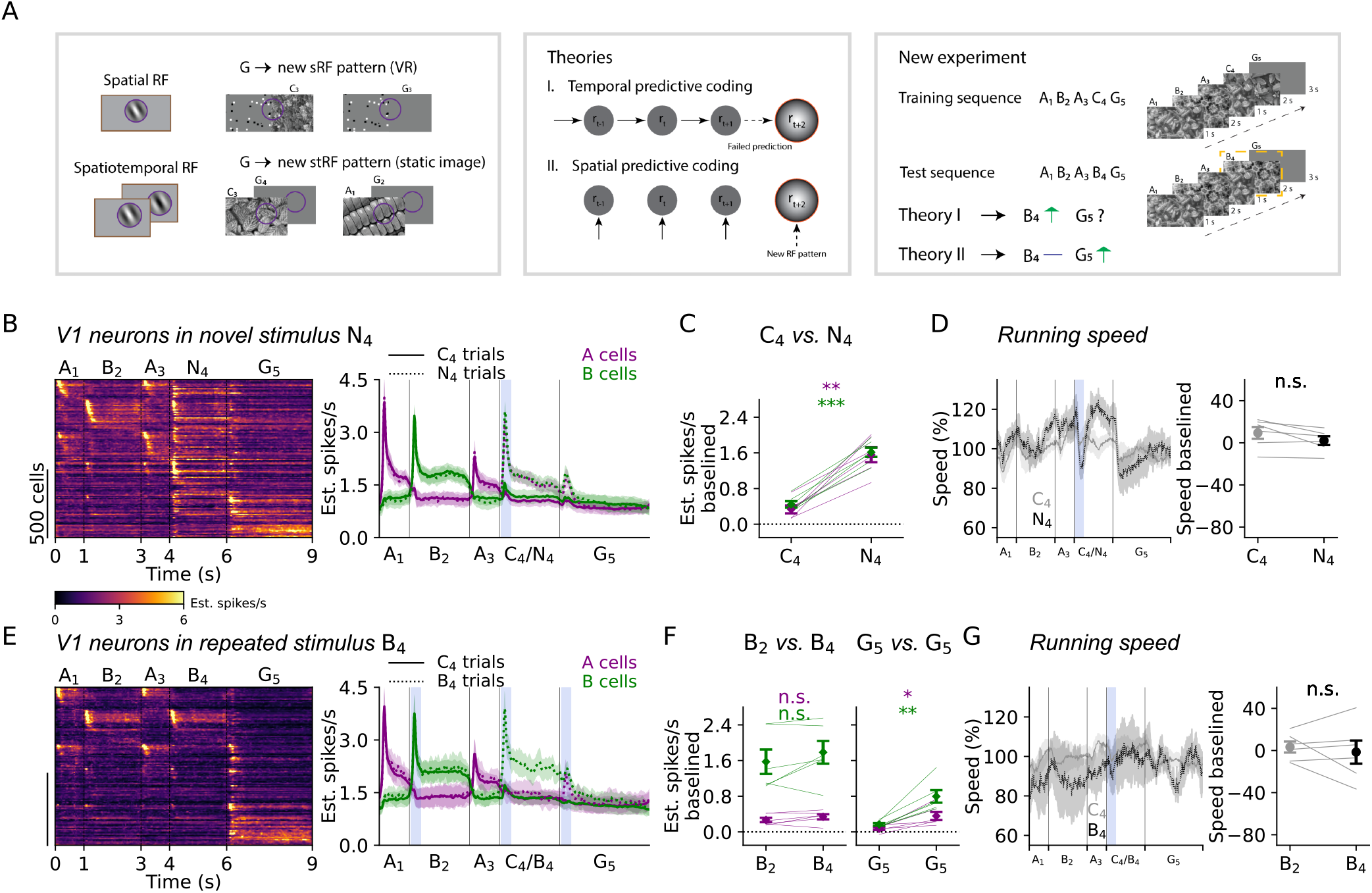
Novel spatial but not temporal patterns drive mismatch responses. (A) (left) Illustration of bottom-up receptive field mismatch in “omission” trials, where an adapted stimulus is replaced with a gray screen. Note that for static images, receptive fields at image transition need to take into account the relative changes from the previous stimulus to the next, similar to how transient retinal cells respond to illumination changes. (middle) Temporal and spatial predictive coding theories suggest different causal relations in the prediction. (right) New experiment design to disentangle the predictions of spatial and temporal predictive coding on static image experiments. (B) (left) Trial-averaged neural responses to A_1_B_2_A_3_N_4_G_5_ trials after training with A_1_B_2_A_3_C_4_G_5_ sequences. Neurons were sorted by their peak response times and held-out trials are shown. (right) Population-average for A cells and B cells (n=6 mice). (C) Quantification of baselined average responses inside the blue window in (B). (D) (left) Average running speeds for A_1_B_2_A_3_C_4_G_5_ and A_1_B_2_A_3_N_4_G_5_ trials, and (right) quantification in the blue shaded area. (E-G) Similar to (B-D) when the C_4_ stimulus is replaced with a B_4_ stimulus (n=6 mice). (F) Comparing responses to (left) B_2_ vs B_4_ in A_1_B_2_A_3_B_4_G_5_ trials or (right) to G_5_ in A_1_B_2_A_3_C_4_G_5_ vs A_1_B_2_A_3_B_4_G_5_ trials. All data are mean*±*s.e.m. across mice. Paired two-sided t-tests were performed across mice.

To further distinguish between the temporal and spatial predictive coding, we designed a new experiment, in which we trained mice on the sequence A_1_B_2_A_3_C_4_G_5_ (Figure 4A). After training, we introduced a test sequence where we replaced C_4_ with B_4_, resulting in A_1_B_2_A_3_B_4_G_5_ trials. Temporal predictive coding would postulate that a prediction error would occur in this case, because the expected transition A_3_ → C_4_ did not happen. The bottom-up or spatial version of predictive coding would not postulate a prediction error, because the transition that is introduced in this way A_3_ → B_4_ was already present in the training stimuli as A_1_ → B_2_. Furthermore, the spatial theory predicts a mismatch error for the transition B_4_ → G_5_, since this was not present in the training stimuli (Figure 4A).

We first verified that under this protocol a mismatch response can indeed still be generated by the introduction of a novel stimulus N_4_ replacing B_4_ (Figure 4B-D). We then focused on trials where C_4_ was replaced with B_4_ (Figure 4E). In these trials, responses to B_4_ were not different from the responses to B_2_, indicating the absence of a prediction error (Figure 4E-G). However, responses to G_5_ on B_4_ trials were greater than on the trained C_4_ trials, in accordance with the prediction of the spatial theory. Thus, elevated neural responses in the static image experiments are also likely due to spatial rather than temporal predictive errors.

### Extended temporal changes drive increased neural activity

Finally, we present a set of experiments that suggest a role for temporal mismatch modulation under more exaggerated manipulations. These experiments were performed using the VR paradigm, and took advantage of the definition of C_3_ as two consecutive “c” images. We first extended the stimulus by another “c”, thus replacing C_3_ with ccC_3_ (Figure 5A). Similar to the experiments with a single “c”, no new spatial patterns were introduced in this way. However, there was slightly elevated neural firing in the extended neural response to the third “c” in the sequence, higher than the response to the second “c”, but similar to the response to the first “c” (Figure 5AB), with no detectable behavioral change (Figure 5C). Next, we repeated the “c” stimulus eight times in a sequence (Figure 5D). For both C and D cells, we observed ramping of neural activity throughout this extended stimulus (Figure 5DE). The response to the last “c” in the sequence was substantially elevated compared to the second “c” (Figure 5E), with no behavioral change (Figure 5F). This ramping response was somewhat surprising, because one may predict adaptation rather than facilitation when the same stimulus is repeated many times. This facilitation was not present in naive animals (Figure 5G-I), ruling out potential accumulation of the jGCaMP8s sensor. We interpret this facilitation as a gain modulation due to a gradual increase in an internal state of alertness or “surprise”. This change in state likely originates elsewhere in the brain, but may lead to differential sensory processing, similar to how other internal states modulate sensory responses [44].

**Figure 5:**
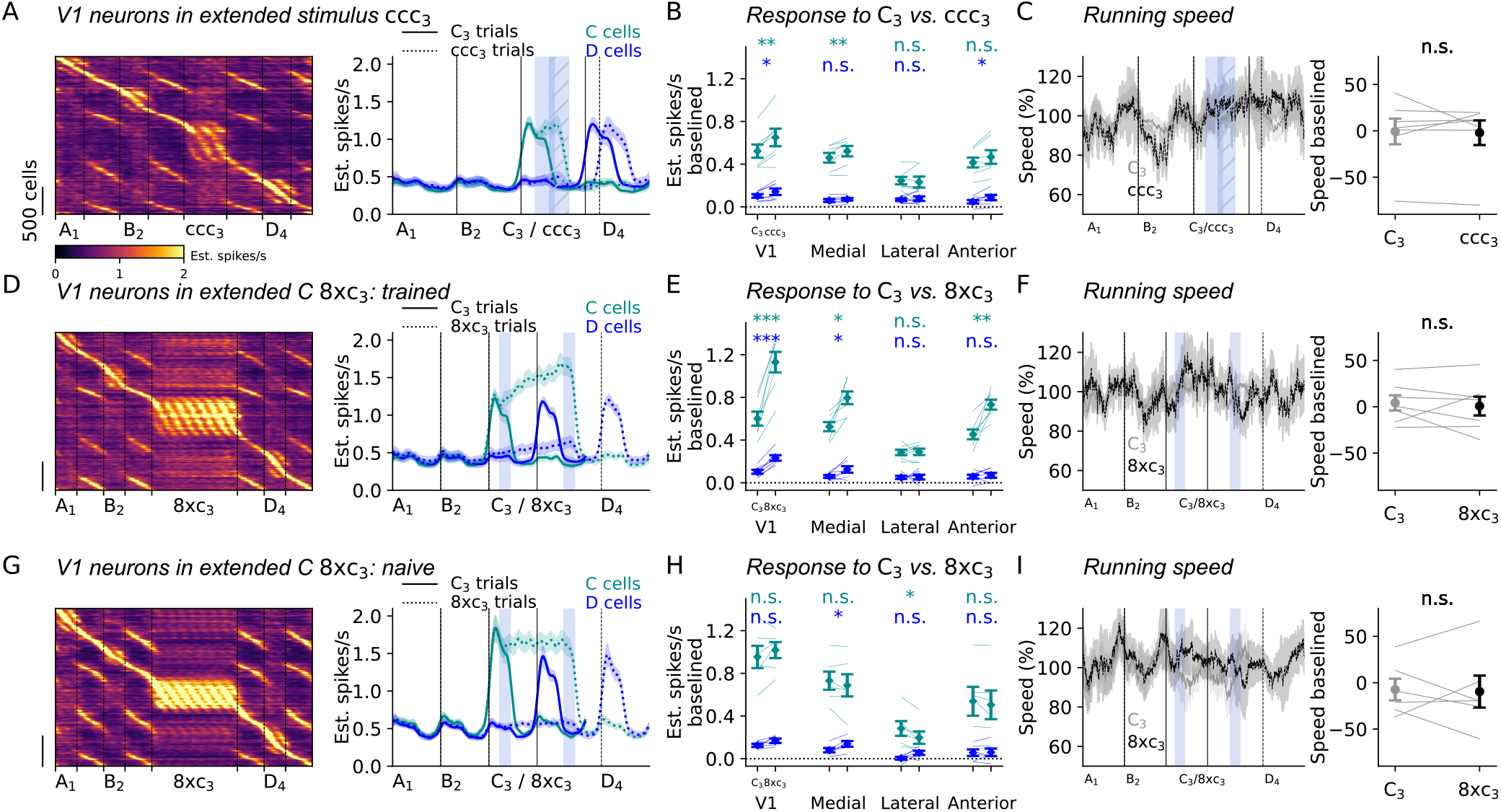
Responses to extended stimuli. (A-C) Same as Figure 1D-F for a test stimulus manipulation where the C = cc stimulus was extended with another copy of c (n=7 mice). Note that this does not create new receptive field patterns (of the kind in Figure 4A). (D-F) Same as (A-C) for 8 consecutive copies of c (n=7 mice). (G-I) Same as (D-F) before training (n=6 mice). All data are mean*±*s.e.m. across mice. Paired two-sided t-tests were performed across mice.

## Discussion

Our results show that continued exposure to a visual stimulus sequence does not result in a temporal mismatch response when that sequence is changed, whether in a closed-loop virtual reality paradigm, or in a closed-loop static stimulus paradigm. Rather, the elevated neural responses signal novel spatial patterns and spatial transitions, previously not viewed during training. Our observations provide evidence to the original formulation of predictive coding as a spatial prediction framework [7].

Other studies have argued against temporal predictive coding in the primary visual cortex of mice, monkeys and humans using oddball paradigms [45, 46]. However, these studies either did not include training over days [45], or did not demonstrate a behavioral or neural effect of training over days [46], leaving open the possibility that the animals did not pay attention to the stimuli and thus could not form an internal model of the sequence. In contrast, we used paradigms that are similar to those in which mismatch responses were previously reported [5, 6, 15]. We demonstrated clear evidence of mismatch to novel spatial patterns, accompanied by behavioral responses during mismatch, and showed that these effects were not present in naive mice. We also used closed-loop experiments, which have previously been suggested to be important for predictive coding [39–42], and we used salient, ecologically-relevant natural images that were not repeated in an oddball paradigm. Briefly, we distinguish here between studies like ours where a stimulus sequence is habituated over many sessions, thus becoming “learned”, and those where sensorimotor coupling itself is being tested with non-”learned” stimuli, such as in virtual reality environments in closed/open loop. We note that under those paradigms too, it is unclear if temporal predictive processing needs to be invoked to explain mismatch responses, because the neural sensory response properties may be sufficient to explain elevated responses [47]. Together with our study, this suggests that temporal prediction is not a fundamental aspect of sensory processing, either in primary or in higher-order sensory areas.

The spatial prediction framework, however, may be an important mechanism by which the brain forms internal models of the visual world. Practical machine learning methods, such as masked autoencoders, demonstrate that a spatial prediction criterion can indeed be leveraged to form such internal models [12]. The biological implementation of such models remains highly unspecified. The mismatch response may be “bottom-up”, for example due to adaptation to familiar images in the feedforward thalamocortical projection, though this is somewhat argued against by the lack of mismatch responses in anesthesia (Figure S2). It could also be “top-down”, such as through a high-level “memory bank” which detects the novelty of a stimulus, and modulates the processing of sensory information in sensory areas through feedback mechanisms [48]. Finally, lateral processing through horizontal connections [49] may implement a version of the masked autoencoder approach, where representations of local image patches are predicted through lateral “attention” mechanisms similar to those in the transformer architecture. This top-down or lateral feedback could be implemented via loops with higher-order thalamic nuclei [15], dendritic modulation [50, 51], and/or specific inhibitory neural targeting [22, 52, 53]. While many possibilities remain open, the precise mechanisms of the spatial pattern learning algorithm remain to be uncovered.

## Acknowledgments

This research was funded by the Howard Hughes Medical Institute at the Janelia Research Campus. From the Vivarium, we thank Jim Cox, Crystall Lopez, Anne Kuzspit, Miriam Rose, Michalis Michaelos, Gillian Harris, Sarah Lindo, Rachel Gattoni and their respective teams for animal breeding, husbandry, surgeries, and behavioral training support. From JeT, we thank Daniel Flickinger, Vasily Goncharov, Alex Sohn, Tobias Goulet, and Steven Sawtelle for help with rig maintenance and upgrades. From MBF Bioscience we thank Georg Jaindl, Mitchell Sandoe, and Boris Djiguemde for scanimage support. We thank Nelson Spruston, Sandro Romani, and Vivek Jayaraman for helpful discussions about the work.

## Author contributions

QZ and MP designed the study. AG performed the surgeries. QZ performed the experiments. QZ performed data analysis with advice from MP and CS. SG, KKL and MF provided gcamp8 viruses. QZ, CS, and MP wrote the manuscript with input from all authors.

## Data availability

The data will be made available upon publication in a journal.

## Code availability

Analysis code will be made available upon publication in a journal.

## Methods

All experimental procedures were conducted according to IACUC, and received ethical approval from the IACUC board at HHMI Janelia Research Campus.

## Experimental methods

### Animals

Eight mice were used in the virtual reality (VR) experiments (4 females, 4 males), ranging in age from 5 to 15 months. Of these, five mice were bred to express GCaMP8s (TetO-jGCaMP8s X CamK2a-tTa, JAX 037717, JAX 007004) and three expressed GCaMP6s (TetO-GCaMP6s × CamK2a-tTa, JAX 024742, JAX 007004) [54]. Nine mice were used in the static image experiments (5 females, 4 males), ranging in age from 4 to 15 months. Of these, five mice were bred to express GCaMP8s (TetO-jGCaMP8s X CamK2a-tTa, JAX 037717, JAX 007004); three transgenic mice (VGAT-CRE x Ai14, JAX 016962, JAX 007914) and one C_57_Bl/6Crl received virus injections to express soma-targeted RiboL1-jGCaMP8s tethered to the ribosome [55]. Mice were housed in reverse light cycle (12 h light on, 12 h light off).

### Surgical procedures

Mice were anesthetized briefly using 3% isoflurane until breathing slowed to approximately 1-2 breaths per second. The animal was transferred to the nosecone on the stereotax. The isoflurane was maintained between 1.5-2% to ensure the animal was within a surgical plane of anesthesia and breathing was approximately 1-2 breaths per second. Marcaine (2.5 mg/kg) was injected subcutaneously beneath the incision area, warmed LRS fluids (+ 5% dextrose) and buprenorphine 0.1 mg/kg (systemic analgesic) were administered subcutaneously along with dexamethasone 7 mg/kg via intramuscular route. Eye lubrication was applied. An incision was made. Vetbond was applied to the skin around the edges of the incision area. Measurements were taken to determine bregma-lambda distance. The center of the right V1 window was measured from lambda: anterior +0.9 mm, lateral +2.6 mm. The edges of the 4 mm craniotomy were measured from this center point, 2 mm in each direction, and the coordinates were lightly etched in the skull using a dental drill. A thin layer of Optibond Solo Plus was applied to the entire surface of the skull and UV cured for 40 s. The headbar was placed over the marked craniotomy location while following the curvature of the skull, then secured using Calibra Universal Resin Cement. The craniotomy was drilled slowly. The skull was also cooled with chilled cortex buffer every few minutes. Once the edges were sufficiently thin, cold sterile-filtered cortex buffer was placed over the well of the headbar and allowed to sit for several minutes. Once the edges softened, the entire bone piece was carefully lifted away using Dumont forceps. A 4+5 mm double window was placed into the craniotomy so that the 4 mm window replaced the previously removed bone piece and the 5 mm window lay over the edge of the bone. Vetbond was applied around the edges of the window. The window was held in place for 5 minutes until the Vetbond fully dried. Then, Calibra Universal Resin Cement was applied over the edges of the 5 mm window and covering the inner sides of the headbar and UV cured. The mouse was removed from the stereotax. Carprofen 5 mg/kg was administered subcutaneously and the animal was allowed to recover on heat. The post-operative care consisted of 2 additional days of monitoring for pain or distress, warm LRS fluids (+5% dextrose), and Carprofen (5 mg/kg) delivered subcutaneously.

For the mice that received viral injections, in addition to the procedures described above, we delivered a 1:1 mixture of Thy1s:TTA (AAV9, 1.64 *×* 1013 V.G./ml) and TRE3G:RiboI1-jGCaMP8s (AAV9, 2.45 *×* 1013 V.G./ml). A total volume of 150 nL was injected across three sites at a depth of –0.3 mm to ensure even expression throughout the cranial window. The injection coordinates relative to lambda were: (1) AP +1.9 mm, ML +2.6 mm; (2) AP 0 mm, ML +1.6 mm; and (3) AP 0 mm, ML +3.6 mm.

### Imaging acquisition

A 2p-RAM mesoscope [56] was used for two-photon imaging. We did online *Z* correction while recording with ScanImage [57]. Recordings for VR experiments were performed at 3.17 Hz and for static image experiments they were performed at 29-30 Hz. Three mice used in the static image experiments were recorded with a Bergamo II microscope at 30 Hz. Data from these animals were combined with those from mice recorded with the mesoscope for subsequent analyses.

During awake recordings, mice were free to run on a styrofoam cylinder. Mice were acclimatized to running on the cylinder in the dark for several sessions before stimulus presentation and imaging.

During anesthetized recordings, mice were briefly anesthetized with 3% isoflurane until respiration decreased to 1–2 breaths/s; isoflurane was subsequently maintained at 1% to stabilize breathing at 1 breath/s. A warming pad was used to maintain body temperature.

### Visual stimuli

The same visual stimulation system was used as in [58]. Three LED tablet screens (aspect ratio 4:3) surrounded the head-fixed mouse covering 270 degrees in azimuth. The mouse was placed in the center of the screen, at a slightly lower elevation from the vertical center.

### VR presentation

The VR environment was rendered with OpenGL in MATLAB. The stimuli were presented with Matlab Psychtoolbox [59]. Natural images A, B, C, D formed the core scene. The images were concatenated, interleaved with random-dot gray walls, and placed on the walls of a linear corridor (Figure 1A). The complete concatenated image contained 2,700 horizontal and 150 vertical pixels. We used pixels as a unit of distance travelled in VR. For closed-loop VR, the VR progressed with the locomotion of the animal: if the mouse speed exceeded a threshold (1.5 cm/s), the corridor advanced at a constant speed (180 pixels/s). For open-loop VR, the VR progressed at a constant speed of 180 pixels/s independent of the mouse running. For the standard A_1_B_2_C_3_D_4_ sequence, it took 15s to finish one trial if the mouse continuously exceeded the running speed threshold.

### Static image presentation

Static images were presented with Matlab Psychtoolbox [59]. Natural images A, B, C, D formed the core scene. In the A_1_B_2_C_3_G_4_ paradigm, stimuli stayed on the screen for 2, 1, 3, 1 seconds, respectively. In the A_1_B_2_A_3_C_4_G_5_ paradigm, stimuli stayed on the screen for 1, 2, 1, 2, 3 seconds, respectively. The progression of stimulus timing was counted only if the mouse was running. Stationary periods were defined as epochs in which the running speed remained below 1 mm/s for at least five consecutive monitor frames (running at 60 Hz). Therefore, if the mouse was stationary, the same stimulus would stay on the screen.

### Behavioral training

Across both the VR and static image paradigms, mice underwent a 1-week training period. During the first 5–7 days they experienced only the standard sequences. Beginning thereafter, we introduced manipulations on a subset of trials.

Before training, we collected a baseline recording that included the standard sequence and all manipulations used later, presented in randomized order. During the post-training test period, we typically ran 1–2 manipulation types per day. Manipulation trials accounted for 7–12% of the total number of trials.

For the VR experiments, we performed a different manipulation on each day, in the following orders over days:

- batch 1 (3 mice): A_1_B_2_D3D_4_, A_1_B_2_G_3_D_4_, A_1_B_2_C_3_D_4_, A_1_B_2_N_3_D_4_, A_1_B_2_ccC_3_D_4_, A_1_B_2_8xC_3_D_4_
- batch 2 (4 mice): A_1_B_2_D3D_4_, A_1_B_2_G_3_D_4_, A_1_B_2_N_3_D_4_, A_1_B_2_C_3_D_4_, A_1_B_2_ccC_3_D_4_, A_1_B_2_8xC_3_D_4_

Two different sets of natural images (A, B, C etc.) were used in those two VR batches. The mice ran 100-300 trials per day with a session duration of 1-1.5 h. If the mouse ran less than 100 trials, the session was excluded from analysis. Some mice stopped running and thus did not complete all manipulation sessions.

For the static image experiments, we did manipulations over days in the following order:

- A_1_B_2_C_3_G_4_ (5 mice): A_1_G_2_C_3_G_4_, A_1_N_2_C_3_G_4_
- A_1_B_2_A_3_C_4_G_5_ (6 mice): A_1_B_2_A_3_B_4_G_5_, A_1_B_2_A_3_N_4_G_5_

Two different sets of natural images were used in these two static-image paradigms. The mice ran 100-400 trials per day with a session duration of 1-1.5 h.

## Data analysis

For analysis we used Python 3 [60], using numpy and scipy [61, 62]. The figures were made using matplotlib and jupyter-notebook [63, 64].

### Processing of calcium imaging data

Calcium raw data were processed with Suite2p [65]. We used Suite2p for motion correction, ROI and neuropil extraction. The neuropil signal was subtracted from the ROI signal with a coefficient of 1. All ROIs with an SNR greater than 0.3 were included for subsequent analysis. The SNR definition was:

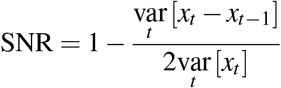

Spike deconvolution was carried out using a new algorithm implemented in Suite2p [65], which employs a neural network trained to predict ground-truth spikes from calcium traces [66].

### VR analysis

We defined trial types based on the manipulation type and location of the manipulation, resulting in the following naming conventions:

- C_3_ trials: A_1_B_2_C_3_D_4_ sequence
- N_3_ trials: A_1_B_2_N_3_D_4_ sequence
- D3 trials: A_1_B_2_D3D_4_ sequence
- G_3_ trials: A_1_B_2_G_3_D_4_ sequence
- C_3_ trials: A_1_B_2_C_3_D_4_ sequence
- 8xC_3_ trials: A_1_B_2_8xC_3_D_4_ sequence

### Train/test split and cell ordering

Normal C_3_ trials were split into training and test sets with 50% of trials in each set. Using the training set, we aligned and sorted cells by the location of their peak response along the linear corridor. This fixed ordering was then applied to the test set (Figure 1C). We defined cell classes by the stimulus window containing their peak. Cells with peaks during the A, B, C, or D image windows were classified as A, B, C, or D cells, respectively, whereas cells with peaks during the random-dot gray periods were classified as G cells.

### Baseline and response windows

Unless otherwise specified, responses for comparisons between standard and manipulated trials were taken from the manipulation window and baselined by subtracting the mean response in a window [−150, −50] pixels preceding the manipulation window. Responses were quantified on the test trials as defined above.

In the second VR batch, We used two novel stimuli in the novelty session and analyzed the data by combining trials from both (N_3_ trials). To compare B and D cell responses between C_3_ and N_3_ trials, we determined the cell classes using training C_3_ trials and assigned cells to the image where their peak firing occurred, excluding the C_3_ image itself. This ensured that neither C_3_ nor N_3_ influenced the cell class assignment, thus allowing for an unbiased comparison between responses to these two stimuli. Responses were computed as the mean activity in the 300 pixel window following C_3_/N_3_ onset (approx. 1,667 ms), after baseline subtraction in the window [−150, −50] before C_3_/N_3_ onset (Figure 1E); running speed and pupil size were evaluated over the same window (Figure 1F, Figure S1A).

Similarly for comparing between D_3_ and D_4_ responses, we sorted the D_3_ trials based on the sorting order from C_3_ train trials without including the C_3_ image responses. Neuronal response within the D_4_ window (300 pixels after stimulus onset, approx. 1,667 ms) from C_3_ trials was compared with that within D_3_ window from D_3_ trials after baseline subtraction (Figure 1H); running speed and pupil size were measured in the corresponding windows (Figure 1I, Figure S1B).

For G_3_ analyses, we again used the C_3_-based ordering without C_3_ responses. We quantified the boundary response when the animal entered the C_3_ image versus G_3_ pure gray screen [-150, 50] around stimulus transition after baseline subtraction (Figure 2EH); running speed and pupil size were compared over the same window (Figure 2FI, Figure S1D).

When analyses included C cells (e.g., C_3_, ccC_3_, 8×C_3_), sorting was derived from C_3_ train trials with C_3_ responses considered. For the shortened stimulus C_3_ condition, the offset response to the second c in C_3_ and the only c in C_3_ (150 pixels after the corresponding stimulus offset, approx. 833 ms) were compared after subtracting the baseline ([*−*150, −50] pixels before the first c onset) response (Figure 2B). For the extended stimulus ccC_3_ condition, the response to the second c and some offset in C_3_ and the third c and some offset in ccC_3_ (200 pixels after the corresponding stimulus onset) were compared after subtracting the baseline ([−150, −50] pixels before the first c onset) response (Figure 5B). For the extended stimulus 8xC_3_ condition, the response to the second c in C_3_ and the last c in 8xC_3_ (−150 pixels after the corresponding stimulus onset) were compared after subtracting the baseline ([−150, 50] pixels before the first c onset) response (Figure 5EH). Running speed and pupil size were evaluated in the corresponding windows.

### Static image analysis

We defined trial types based on the manipulation type and location of the manipulation, resulting in the following naming conventions:

- B_2_ trials: A_1_B_2_C_3_G_4_ sequence
- N_2_ trials: A_1_N_2_C_3_G_4_ sequence
- G_2_ trials: A_1_G_2_C_3_G_4_ sequence
- C_4_ trials: A_1_B_2_A_3_C_4_G_5_ sequence
- B_4_ trials: A_1_B_2_A_3_B_4_G_5_ sequence
- N_4_ trials: A_1_B_2_A_3_N_4_G_5_ sequence

Cells were sorted in the same way as in the VR experiment. The normal trials (either A_1_B_2_C_3_G_4_ or A_1_B_2_A_3_C_4_G_5_ sequence) were split into training and test data. We aligned cells based on their peak response time in the static image sequence within the training data. This sorting order was applied to the test data (Figure 3B). We defined cells by their peak response position in the A_1_ stimulus window as the A cells (and similarly for B, C, G cells).

We used two novel stimuli in the novelty session and analyzed the data by combining trials from both in the N_2_ and N_4_ trial analyses.

For the A_1_B_2_C_3_G_4_ sequence, to compare the responses of A, C cells in B_2_ and N_2_ (or G_2_) trials, in the training set with the B_2_ trials, we aligned cells without using B_2_ image responses. Neuronal response within the B_2_/N_2_ (or G_2_) window (0-300 ms after stimulus onset) was averaged and baselined using the final 100 ms of the gray screen period (Figure 3DG). When comparing the running speed in B_2_ and N_2_ (or G_2_) trials, traces were extracted from 0-400 ms after stimulus onset and baselined by the final 200 ms of the gray screen period.

Similarly, for the A_1_B_2_A_3_C_4_G_5_ sequence, to compare the responses of A, B cells in C_4_ and N_4_ trials, responses were extracted from C_4_ and N_4_ window (0-300 ms after stimulus onset) and baselined with the final 100 ms of the gray screen period (Figure 4C). For the A_1_B_2_A_3_B_4_G_5_ sequence, when comparing responses in B_2_ and B_4_, responses were extracted from B_2_ and B_4_ window (0-300 ms after stimulus onset) and baselined using the final 100 ms of the gray screen period of B_4_ trials (Figure 4F, left); when comparing responses in G_5_ from C_4_ and B_4_ trials, responses were extracted from G_5_ window (0-300 ms after the onset of gray screen) and baselined (the final 100 ms of the gray screen period) of C_4_ and B_4_ trials, respectively (Figure 4F, right). The running speed during B_2_ and B_4_ stimuli in B_4_ trials was defined as the average running speed 0-400 ms after stimulus onset, and was baselined by the final 200 ms of the gray screen period (Figure 4G).

### Statistical tests

All the statistics were paired two-sided t-tests across mice. Detailed *P* values are reported below.

- 1E. B cells: 2.68*×*10*−*4 / 1.41*×*10*−*3 / 9.17*×*10*−*3 / 1.76*×*10*−*3; D cells: 4.83*×*10*−*4 / 4.96*×*10*−*3 / 5.90*×*10*−*3 / 0.0731
- 1F. 0.0348
- 1H. B cells: 0.370 / 0.0815 / 0.478 / 0.599; D cells: 0.348 / 0.165 / 0.599 / 0.920
- 1I. 0.249
- 1K. B cells: 2.54 *×*10*−*3 / 0.0990 / 0.107 / 0.116; D cells: 0.0419 / 0.0440 / 0.499/ 0.0723
- 2B. C cells: 0.133 / 0.184 / 0.827 / 0.586; D cells: 0.0588 / 0.121 / 0.861 / 0.0742
- 2C. 0.199
- 2E. B cells: 0.685 / 0.0543 / 0.0574 / 0.0365; D cells: 5.30*×*10*−*3 / 0.0191 / 0.0162 / 0.0713; G cells: 3.27*×*10*−*3 / 9.49*×*10*−*3 / 0.0891 / 0.105
- 2F. 0.0179
- 2H. B cells: 5.08 *×*10*−*3 / 0.538 / 0.168 / 0.188; D cells: 0.0281 / 0.431 / 0.872 / 0.182; G cells: 0.0284 / 2.58*×*10*−*3 / 0.0649 / 0.510
- 2I. 0.721
- 3D. A cells: 0.0250; C cells: 8.88*×*10*−*3
- 3E. 0.497
- 3G. A cells: 7.79*×*10*−*3; C cells: 4.94*×*10*−*3
- 3H. 0.312
- 4C. A cells: 1.23*×*10*−*3; B cells: 5.42*×*10*−*5
- 4D. 0.078
- 4F. (left) A cells: 0.174; B cells: 0.214; (right) A cells: 0.0245; B cells: 3.42*×*10*−*3
- 4G. 0.297
- 5B. C cells: 3.40 *×* 10*−*3 / 8.26 *×* 10*−*3 / 0.623 / 0.0948; D cells: 0.0489 / 0.0986 / 0.618 / 0.0335
- 5C. 0.425 5E. C cells: 5.80 *×* 10*−*4 / 0.0126 / 0.838 / 3.00 *×* 10*−*3; D cells: 1.82 *×* 10*−*4 / 0.0437 / 0.934 / 0.548
- 5F. 0.649
- 5H. C cells: 0.282 / 0.181 / 0.0199 / 0.257; D cells: 0.146 / 0.0109 / 0.0965 / 0.852
- 5I. 0.864
- 1S1C. B cells: 0.255 / 0.245 / 0.0690 / 0.760; D cells: 0.637 / 0.203 / 0.839 / 0.193
- 1S2. 0.0237 / 0.730 / 0.235 / 4.50 *×* 10*−*3 / 0.268 / 0.174

### Retinotopy

We used the same procedure as in [58] to estimate spatial receptive locations. A set of 500 natural images, each presented three times, was used for mapping. First, responses from a reference mouse were fitted with 200 spatial kernels. Recordings from all other mice were then fitted using the same model to determine the preferred kernel and corresponding spatial location of each cell. These spatial locations were aligned to the spatial map of the reference mouse, in which the recorded region was segmented according to the computed sign map [67].

---

**Figure S1:**
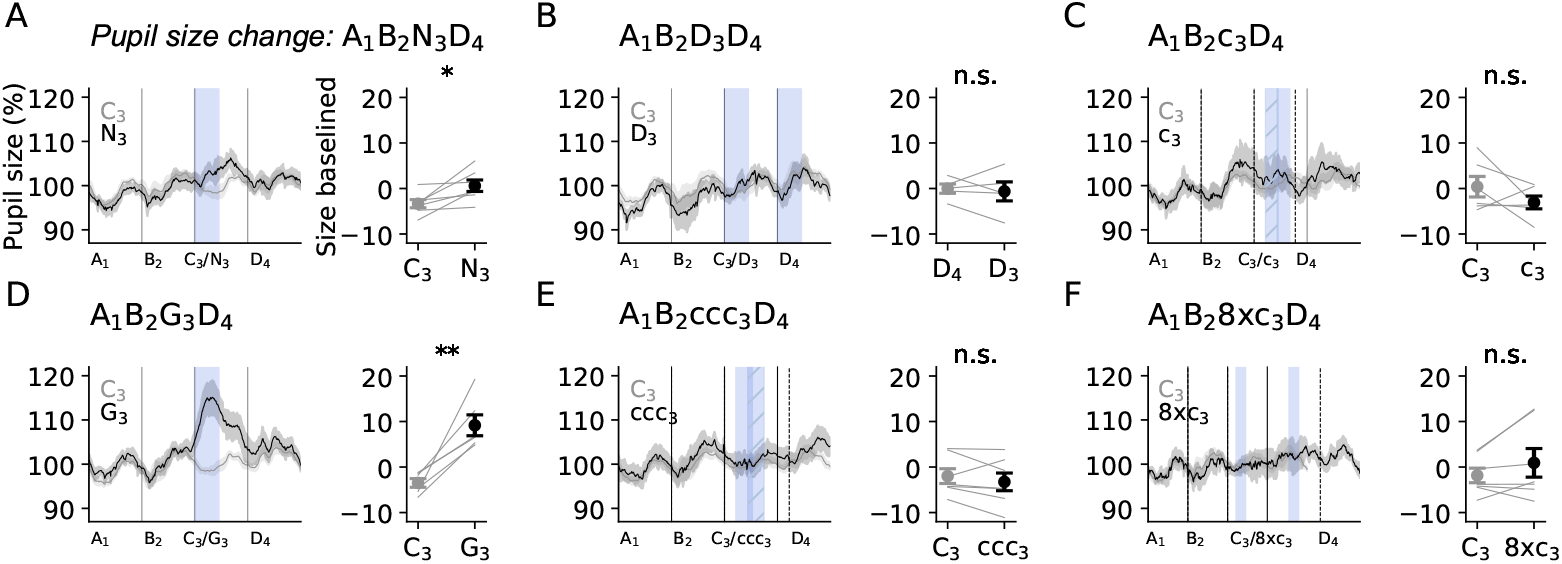
Pupil fluctuations. (A) (left) Average pupil size over trials. (right) Average pupil size in the new stimulus time window compared between C_3_ and N_3_ trials (n=7 mice). (B-F) Same as (A) for test sessions where different stimuli are introduced: (B): D_3_, D stimulus at position 3 (n=5 mice), (C): G_3_, gray screen at position 3 (n=6 mice), (D): c_3_, half of C stimulus at position 3 (n=6 mice), (E): ccc_3_, repeated half of C stimulus at position 3 (n=7 mice), (F): 8xc_3_, 8 times repeated half of C stimulus at position 3 (n=7 mice). All data are mean *±* s.e.m. across mice. Paired two-sided t-tests were performed across mice.

**Figure S2:**
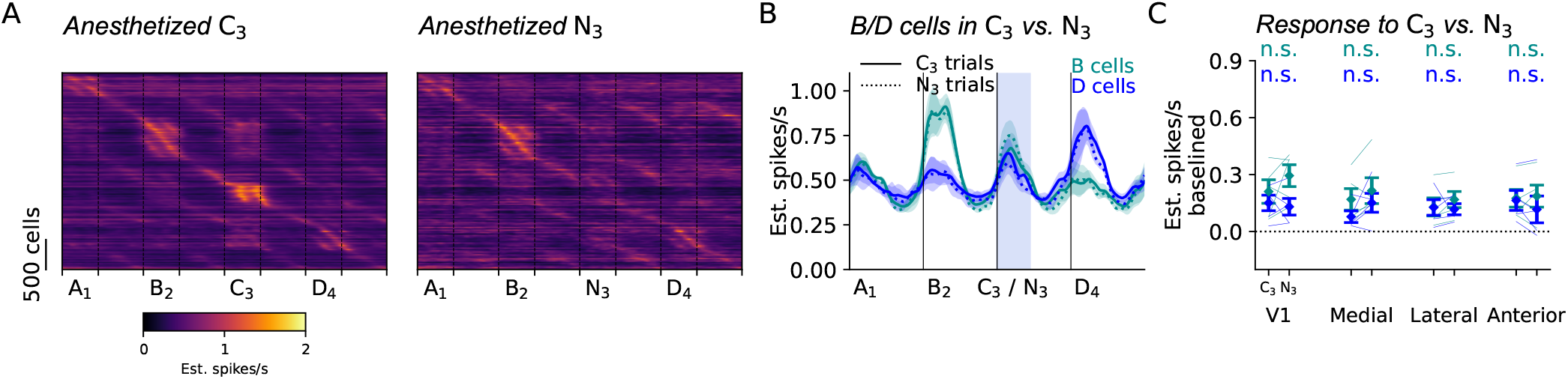
Responses to novel stimuli in anesthesia. (A) Sorted neural responses to the trained stimulus (C_3_) and the novel stimulus (N_3_). (B) Average responses of B and D cells on C_3_ and N_3_ trials (n=4 mice). (C) Average responses during the C_3_ and N_3_ stimulus presentations compared between B and D cells. All data are mean*±*s.e.m. across mice. Paired two-sided t-tests were performed across mice.

